# HeteroRepur: Efficient Modeling of Heterogeneous Disease Graph for Depression Drug Repurposing using Heterogeneous Graph Neural Networks

**DOI:** 10.1101/2024.12.18.629087

**Authors:** Ko-Hong Lin, Tongtong Huang, Kai Zhang, Xiaoqian Jiang, Yejin Kim

## Abstract

**Objective:** Major Depressive Disorder (MDD) is one of the most prevalent psychiatric disorders, yet existing treatments often result in suboptimal responses and encounter high recurrence rate. Repositioning existing medications, which often significantly reduces the required time and budget, has been demonstrated to be a promising strategy for drug discovery of multiple central nervous system (CNS) disorders. However, an efficient modeling strategy of heterogeneous semantic information among biological networks has yet to be developed for drug repurposing.

**Material and Methods:** In this study, we proposed HeteroRepur, by integrating heterogeneous graph neural networks to learn interaction features of Depression Drug Repurposing Graph (DDRG) and then classify drugs.

**Results:** The DDRG embeddings learned by HeteroRepur accurately capture graph information, including 89 types of interaction or associations among drugs, genes, pathways, and GOs, as well as disease genetic network of MDD. In drug classification task, HeteroRepur outperforms various baseline models, including existing graph neural network approaches that do not consider heterogeneous network information. Finally, HeteroRepur predicts new candidates for treating MDD, including CNS drugs that have the potential to be extended to MDD, and dietary supplements that might benefit depression patients by elevating essential nutrient levels.

**Conclusion:** HeteroRepur outperforms traditional graph-based approaches, and can be used to rank promising drugs among complex disease networks with heterogeneous topology.

## 1 Introduction

Major Depressive Disorder (MDD) is one of the most prevalent psychiatric diseases, affecting 280 million people and causing over 700,000 suicidal mortalities [1]. Although many antidepressant drugs have been explored since the first use of Iproniazid in treating MDD in 1958, approximately two-thirds of MDD patients have shown insufficient response to first-line medications [2]. Additionally, 20-30% of MDD patients developed treatment-resistant depression (TRD), failing to respond to at least two antidepressants [3]. These challenges of MDD treatment strategies highlighted the importance of continuous efforts in exploring efficacious medications for MDD.

Alternative to the traditional drug discovery process, drug repurposing is a promising strategy that might bypass the lengthy processes that have already been completed [4]. Drug repurposing in CNS disorders has been especially successful, with around 13 drugs repurposed for treating depression or bipolar depression, and 20 antidepressant treatments have been repurposed for new indications [5]. Given the huge unmet need in the therapeutic area of MDD, the development of computational approaches might help efficiently identify repurposable candidates.

Recent efforts of computational drug repurposing relied on either drug signature matching [6,7], molecular docking [8], genome-wide association studies [9], or network analysis [10]. Each perspective takes advantage of various biological and clinical data (e.g., drug-perturbed gene expression, chemical structure, etc.), and integrating these domains provides complementary knowledge for drug repurposing [11]. Additionally, genetic regulation of MDD are complicated due to the dynamic interplays across the periphery and central nervous system. The etiology of depression is associated with biological responses across serotonergic, glutamatergic, and GABAergic systems, underlying the importance of considering a group of disease-underlying genes rather than a few targets [12]. To integrate these knowledge, the utilization of network-based methodologies can capture the intricate interplays between drugs, proteins, and diseases while incorporating disease modules that represent canonical disease gene subnetworks [13–15]. This is imperative as pharmacological effects not only treat diseases by directly targeting specific proteins but also through the regulation of distant proteins in the same biological functions or pathways [16–18]. Consequently, network-based drug discovery approaches have incorporated hierarchical relationships between gene ontologies and biological pathways [15,19]. Furthermore, chemical structural properties can also functionally impact its treatment efficacy and bioavailability. Collectively, the existing knowledge of above-mentioned biomedical interactions can be integrated into heterogeneous knowledge graphs, which contain multiple types of entities and relationships, for drug repurposing applications [20–22].

Previous studies on drug repurposing using data-driven approaches have attempted to model the complex relationships between biological entities to prioritize drug candidates for repurposing using geometric approaches [23], matrix/tensor factorization [24,25], and graph neural networks (GNNs) [26,27]. However, these approaches often ignore the different interactions or associations between the same entities, and cannot effectively learn to encode distinct semantic information in complex biomedical networks. For example, a drug’s ability to upregulate and downregulate gene expression may elicit opposing effects [28], while a compound inhibiting phosphorylation or promoting ubiquitination of a protein can also lead to diverse responses [29]. These distinct interaction or association types can make varying contributions to the downstream task. Therefore, several heterogeneous graph neural networks [30–32], which consist of certain attention mechanisms that allow efficient modeling of these contrasting relationships, could address the challenges of heterogeneous graph representation learning in biomedical networks.

In this study, we present HeteroRepur, an MDD drug repurposing pipeline on multiple sources of biological interactions using heterogeneous graph neural networks. We built the Depression Drug Repurposing Graph (DDRG), a comprehensive knowledge graph with 89 types of relationships among biological interactome, chemical-chemical association networks, and disease genetics information. The main contribution of this study includes:

1. Designed a comprehensive knowledge graph for MDD drug repurposing by integrating disease genetic interactions, biological networks, and cheminformatics.
2. Developed a heterogeneous graph neural network-based drug repurposing appraoch that outperformed traditional methods which did not consider distinct biological interactions and responses.
3. Our novel predictions include CNS drugs and dietary supplements, which have been reported repurposing potentials for treating depression-like symptoms or alleviating nutrient deficiency in MDD patients [33–35].

By applying HeteroRepur, we explore therapeutic benefits through rich semantic information from literature-based evidence, broadening the scope of drug repurposing that is usually restricted by the limited amount of chemical and genetic data.

## 2 Results

### 2.1 DDRG graph representation learning and applications in drug classification

To embed DDRG features, we first performed graph representation learning, with multiple hyperparameters considered (**Table 1**, Graph representation model). Implementing dropout did not improve link prediction performance. Among 89 DDRG edge types, the graph representation model can achieve AUROC and AUPRC of 0.94 and 0.85 on the least performing edge type in DDRG, demonstrating that multimodal interactions and gene class features were well represented by the embeddings (**Figure 2**).

**Table 1.** Hyperparameters considered through grid search.

| Hyperparameter | Value considered |
| --- | --- |
| Graph representation model |  |
| Learning rate | 0.01; **0.005**; 0.001; 0.0005 |
| Edge sampling ratio | **1:15**; 1:100 |
| Drug classification model |  |
| Loss function | **Focal loss ($\alpha = 1$, $\gamma = 2$)**, Focal loss ($\alpha = 0.75$, $\gamma = 2$), Focal loss ($\alpha = 0.25$, $\gamma = 2$), Focal loss ($\alpha = 1$, $\gamma = 3$), Focal loss ($\alpha = 0.75$, $\gamma = 3$), Focal loss ($\alpha = 0.25$, $\gamma = 3$), Binary Cross Entropy Loss |
| Learning rate | 0.01; 0.005; **0.001**; 0.0005 |
| Dropout | **0**; 0.5 |

To predict MDD drugs using the DDRG embeddings, HeteroRepur utilized an RGCN drug classification model that classifies positive and negative drugs among 1,216 drug candidates. Drug candidates in DDRG were split by 60%, 20%, and 20% for the training, evaluation, and testing stage. We considered several sets of hyperparameters for model optimization (**Table 1**, Drug classification model). Our model can accurately identify positive drugs for treating MDD in DDRG (recall=0.91, accuracy=0.96, precision@50 = 0.98, precision@60 = 0.87).

### 2.2 RGCN-based drug classification model outperforms baseline models

We compared our RGCN-based drug classification model with logistic regression, SVM, XGBoost, and random forest. All of them were trained using DDRG embeddings as input drug features. To compare the performance of each model, we utilized AUROC and AUPRC to evaluate the drug classification on the test set through 5-fold cross-validation (**Table 2**).

**Table 2.**
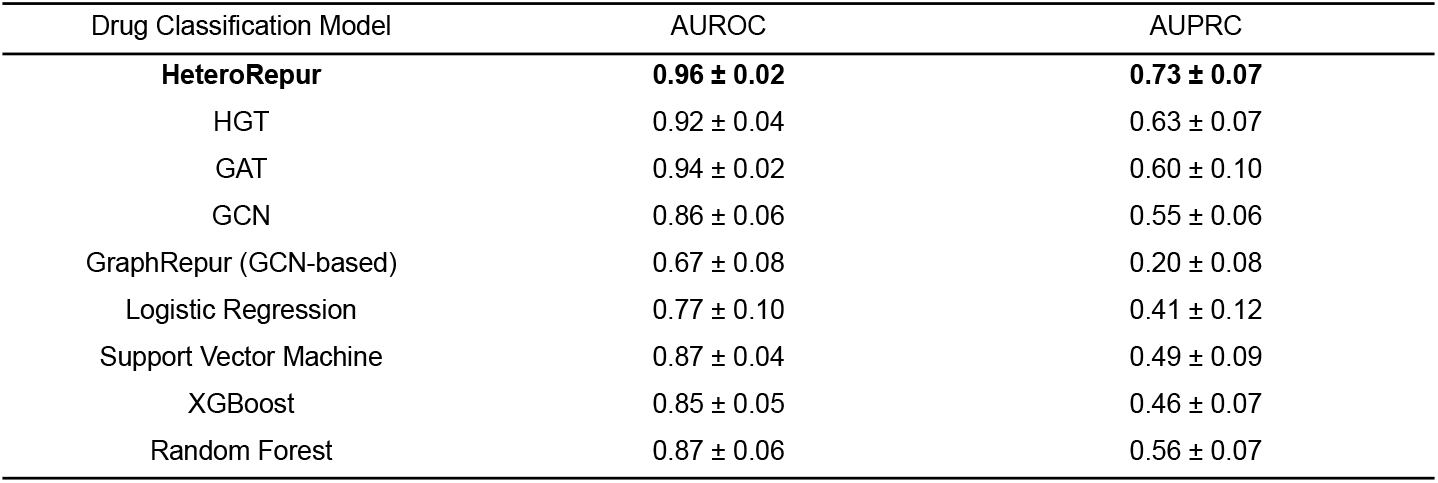
Drug classification model comparisons based on AUROC and AUPRC.

| Drug Classification Model | AUROC | AUPRC |
| --- | --- | --- |
| **HeteroRepur** | **0.96 ± 0.02** | **0.73 ± 0.07** |
| HGT | 0.92 ± 0.04 | 0.63 ± 0.07 |
| GAT | 0.94 ± 0.02 | 0.60 ± 0.10 |
| GCN | 0.86 ± 0.06 | 0.55 ± 0.06 |
| GraphRepur (GCN-based) | 0.67 ± 0.08 | 0.20 ± 0.08 |
| Logistic Regression | 0.77 ± 0.10 | 0.41 ± 0.12 |
| Support Vector Machine | 0.87 ± 0.04 | 0.49 ± 0.09 |
| XGBoost | 0.85 ± 0.05 | 0.46 ± 0.07 |
| Random Forest | 0.87 ± 0.06 | 0.56 ± 0.07 |

Among these models, RGCN outperformed other heterogeneous graph neural networks (Heterogeneous Graph Transformer), homogeneous graph neural networks (Graph Convolution Network, Graph Attention Network), and machine learning models (logistic regression, support vector machine, XGBoost, and random forest). GraphRepur is a GCN-based cancer drug classification model across sparse drug-drug connections, which prevents effective information propagation and limits its performance in drug classification.

### 2.3 Drug-drug associations are a critical component of DDRG for the drug classification model

Next, to understand each edge type’s contribution to predicting positive drugs, we evaluated which edge information of DDRG is the most informative in the drug classification task. Perturbation experiments on different edge types were conducted to compare the changes in the performance of our drug classification model. When perturbing drug-drug associations in both the pre-training step and drug classification task, the AUPRC decreased significantly (∼10%), while ablating GO- and pathway-related edges also slightly reduced the AUPRC values (**Table 3A**). Perturbing each component only in either the pre-training step or drug classification task would not impact the drug classification performance. Finally, substituting the DDRG topology with a drug-drug network would significantly decrease the drug classification performance (**Table 3A**).

**Table 3.** Drug classification performance using different graph components or without pre-training step. (A) Ablation study of DDRG components. (B) Removing pre-training step from the framework.

| (A) |  |  |
| --- | --- | --- |
| Pre-trained Features | Network | AUPRC |
| **DDRG** | **DDRG** | **0.73±0.07** |
| DDRG remove Drug-Drug | DDRG remove Drug-Drug | 0.63±0.13 |
| DDRG remove GOs | DDRG remove GOs | 0.71±0.10 |
| DDRG remove Pathways | DDRG remove Pathways | 0.67±0.09 |
| DDRG | Drug-drug network | 0.24±0.05 |
| (B) |  |  |
| Pre-trained Features | Network | AUPRC |
| Random | DDRG | 0.66±0.09 |
| Random | DDRG w/o drug-drug | 0.60±0.09 |
| Random | Drug-drug network | 0.18±0.07 |

To study the importance of the pre-training step in the framework, we used random features and trained the drug classification model only through the DDRG topology (**Table 3B**). The hyperparameters of all models were obtained from the best performance in **Table 1**. With 5-fold cross-validation, we demonstrated that the pre-training step can essentially improve the drug classification model performance in all types of network structures (DDRG, DDRG without drug-drug associations, and drug-drug network only).

### 2.4 Novel Drug Prediction

Model hyperparameters of the best-trained model were used to test the complete set of drugs in DDRG. Since the positive set already includes FDA-approved drugs, investigational drugs currently under clinical trials, and drugs with potential therapeutic roles, our goal is to examine if our model could identify additional promising drugs. Among the top 100 drugs, 59 drugs are approved medications, investigational drugs, or therapeutically promising for MDD listed in the positive set. Novel predictions of drugs that have supportive evidence of beneficial effects for MDD are listed in **Table 4**.

**Table 4.** Novel predictions of our study.

| Drug name | Original Indication(s) | Supporting evidence |
| --- | --- | --- |
| MDMA | N/A | [59] |
| Magnesium | Hypomagnesemia; Seizure | [60,61] |
| Ondansetron | Postoperative Nausea; Vomiting | [47,62] |
| Buspirone | Anxiety Disorders | [55,56] |
| Vitamin E | Vitamin E Deficiency | [63,64] |
| Caffeine | Apnea; Bronchopulmonary Dysplasia | [48,49] |
| Vitamin D | Hypoparathyroidism; Refractory Rickets;<br>Familial Hypophosphatemia | [34,57] |
| Coenzyme Q10 | Congestive Heart Failure; Systolic<br>Hypertension | [35,58] |
| Bosentan | Pulmonary Arterial Hypertension | [50] |

Among the novel predictions, many of them have been demonstrated to play beneficial roles in depression or other psychiatric disorders. Ondansetron is a 5-hydroxytryptamine 3 (5-HT3) receptor antagonist (5-HT3RAs) approved for treating Nausea and vomiting. [47] have shown the in vivo efficacy of Ondansetron in attenuating depression and anxiety associated with obesity by inhibiting biochemical alterations and improving serotonergic neurotransmission. It is currently undergoing a phase 4 trial for treating MDD (ClinicalTrials.gov Identifier: NCT02082678). A cross-sectional and prospective study determined associations between coffee consumption and a significantly lower risk of depression [48] [49]. Bosentan was indicated to treat pulmonary arterial hypertension by targeting Endothelin-1 (ET1), while in vivo evidence has been reported to demonstrate its anti-depressant-like activity, which is possibly associated with increased circulating IL-6 levels [50]. MDMA is a monoamine release and reuptake inhibitor and can enhance synaptic levels of serotonin, norepinephrine, and dopamine [51]. Despite not being tested in clinical trials for treating depression, MDMA has shown beneficial effects as an adjunct to psychotherapy, and significant relief has been reported in an earlier study [52]. Recently, MDMA also showed preliminary but promising effects in post-traumatic stress disorder (PTSD) [53] and social anxiety in autistic adults [54], which are promising to be extended to MDD. Buspirone is efficacious and well-tolerated in the treatment of Generalized anxiety disorder (GAD) in elderly patients [55]. Six weeks of Buspirone and Melatonin combination treatment was found to have significant antidepressant effects [56].

Interestingly, nutrient supplements constitute a pivotal component of our prediction. In a recent randomized controlled trial (ClinicalTrials.gov Identifier: NCT02466087), mild-to-moderate MDD patients receiving magnesium daily for 6 weeks had significant improvement in Generalized Anxiety Disorders-7 scores and Patient Health Questionnaire-9 (PHQ-9) scores [33]. Additionally, Magnesium supplements were well tolerated by the patients without the need for close monitoring for toxicity. Vitamin E has various body-supporting effects, including anti-inflammatory and antioxidant abilities. However, meta-analyses showed inconclusive results in reducing depression, and future studies will be required with a larger sample size to elucidate the associations between Vitamin E consumption and treating depression. Vitamin D is vital in supporting immune function, neuromuscular function, and reducing inflammation. Increased circulating 25(OH)D concentrations following 8-week vitamin D supplementation (50,000 IU 2 weeks-1) resulted in a significant decrease in BDI-II scores in patients with mild to moderate depression [34]. Umbrella meta-analysis also confirms the potential benefits of vitamin D supplementation and higher serum vitamin D levels in reducing the development and symptoms of depression [57]. An in vivo study found that Coenzyme Q10 can reverse streptozotocin-induced depression-like behavior in adult male mice [58]. [35] found that Coenzyme Q10 is a marker for treatment resistance in depression, as a lower CoQ10 level is especially a risk factor for depression in coronary artery disease and chronic heart failure (CHF) patients who took statin medications.

## 3 Discussion

Despite graph neural networks could encode graph structure and transform edge-based knowledge for many drug development tasks, some still suffer from significantly increased amount hyperparameters and scalability problems, as well as difficulties in distinguishing various types of biological interactions. Heterogeneous graph neural networks were designed to efficiently handle heterogeneous graphs and address the challenge by capturing different distributions and characteristics of various edge types between drugs, proteins, and other entities.

In this study, we formulated the MDD drug repurposing task into a drug classification problem, based on drug embeddings of DDRG. HeteroRepur first learned all embeddings in DDRG by capturing heterogeneous interactions and genetic classes, utilizing a Heterogeneous Graph Transformer (HGT), which can perform scalable and efficient training of 89 types of link information [31]. In the drug classification task, the Relational Graph Convolutional Network (RGCN) model [30] outperformed all other baseline models in predicting positive drugs from all drugs among DDRG. With scalability and attention-based mechanisms, deploying heterogeneous graph neural networks in graph representation learning can efficiently embed a large number of interactions in DDRG and also outperform other graph-based models in drug classification.

Our results implied that considering interactions between drugs and proteins, along with pathways, biological function, and disease-specific gene class information, can be informative in identifying positive drugs in MDD. Additionally, pre-training of graph DDRG representation improved performance in predicting positive drugs, while the ablation study showed that each interaction type in DDRG, especially drug-drug associations, is informative in MDD drug prediction. Finally, our novel drug prediction identified drugs that have beneficial roles in depression and other psychiatric disorders, while clinical trials and literature evidence also support several nutrient supplements.

Computational modeling and prediction of drugs were often limited by the accessibility or small sample sizes of public domain data, such as chemical 3D structures, drug perturbation gene signatures, or patient multi-omics data. HeteroRepur integrates existing disease knowledge, therapeutic mechanisms, and biological interactions to prioritize drug candidates, enabling the reevaluation of current medications and compounds for their repurposing potential for Major Depressive Disorder (MDD), without requiring the availability of these data. HeteroRepur also allowed additional opportunities to find benefits in nutrient supplements, which were not included in such datasets. Although the treatment effects of these candidates were less established since fewer clinical trials were designed to study the efficacy of dietary supplements, they usually have better tolerance and marginal side effects. Additionally, there is almost no need for close monitoring for toxicity.

## 4 Conclusion

In this study, we proposed HeteroRepur, based on the integration of heterogeneous graph neural networks and multi-task learning techniques. HeteroRepur learns to represent heterogeneous network features from DDRG and predicts positive drugs for MDD. In comparative studies with baselines, we found that heterogeneous graph neural networks consider distinct semantic information of interactions between drugs and proteins, as well as GO and pathway relationships, outperforming all the homogenous GNNs and machine learning models that do not distinguish heterogeneous edge information. Additionally, we systematically compared the contributions of different edge types and features, demonstrating that distinct edge types are informative in predicting positive drugs. Our analysis showed that HeteroRepur accurately learns distinct DDRG edge information and robustly outperforms other graph-based models in predicting positive drugs. Collectively, our study showcased the advantage of applying scalable and efficient heterogeneous graph neural networks to enhance the accuracy of network-based drug repurposing from complex disease knowledge graphs.

## 5 Materials

### 5.1 Graph Construction

We developed the Depression Drug Repurposing Graph (DDRG), a bi-directional heterogeneous graph that depicts molecular interactions (drug-gene, gene-gene interactions, gene-GO, and gene-pathway associations), chemical structural associations (drug-drug associations), and disease genomics features (gene classes), related to Major Depressive Disorder. The relationships between drugs, genes, GOs, and pathways were obtained from the Comparative Toxicogenomics Database (CTD), by querying MeSH ID “D003865” (Depressive Disorder, Major). Specifically, curated semantic information of drug-protein interactions (E.g., decreases_expression, increases_methylation, affects_binding, etc.) was retrieved and set as distinct edge types. In total, 367,300 edges across 89 edge types with at least 20 interactions were collected. The graph comprises 1,216 unique drugs, 13,689 genes, 2,267 GOs, and 96 pathways. Details of DDRG are shown in **Figure 1a**.

**Figure 1.**
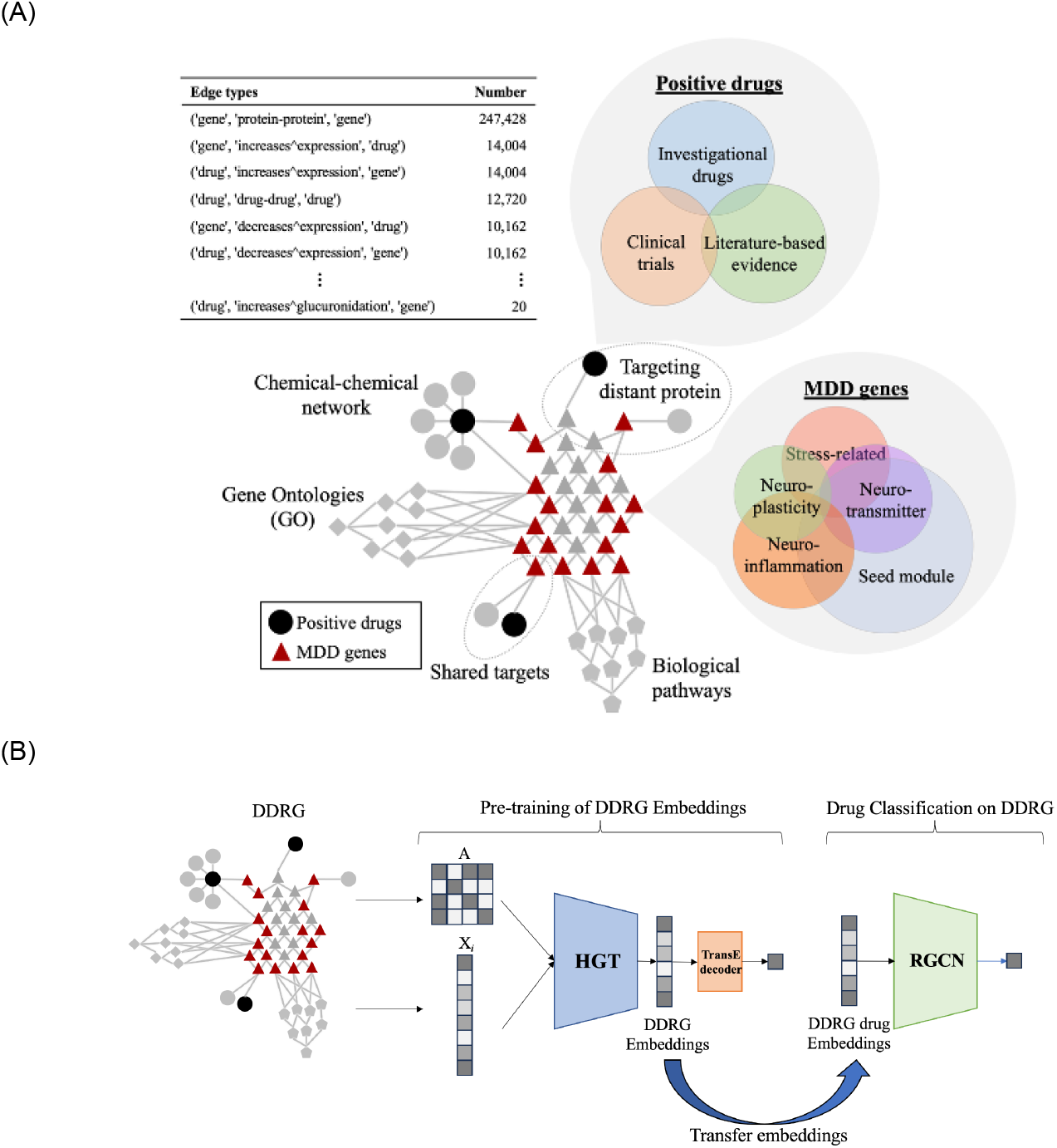
(A) Components of DDRG. DDRG is a bidirectional graph that consists of 89 types of interactions between drugs, genes, pathways, and gene ontologies. Additionally, DDRG includes MDD genes based on previously reported genetic relevance to five major MDD mechanisms. Three groups of positive drugs in DDRG were used in drug classification task. **(B) Schematic representation of model architecture**. Graph adjacency A represents a binary matrix of edge information, while X_*i*_ denotes the input feature of each node *i*. To pre-train DDRG features, an HGT encoder was used to encode the input node feature and edge information, and then a TransE decoder output a TransE score for each edge. After pre-training, DDRG drug embeddings were used for drug classification, which includes an RGCN model that classified positive drugs for MDD.

**Figure 2.**
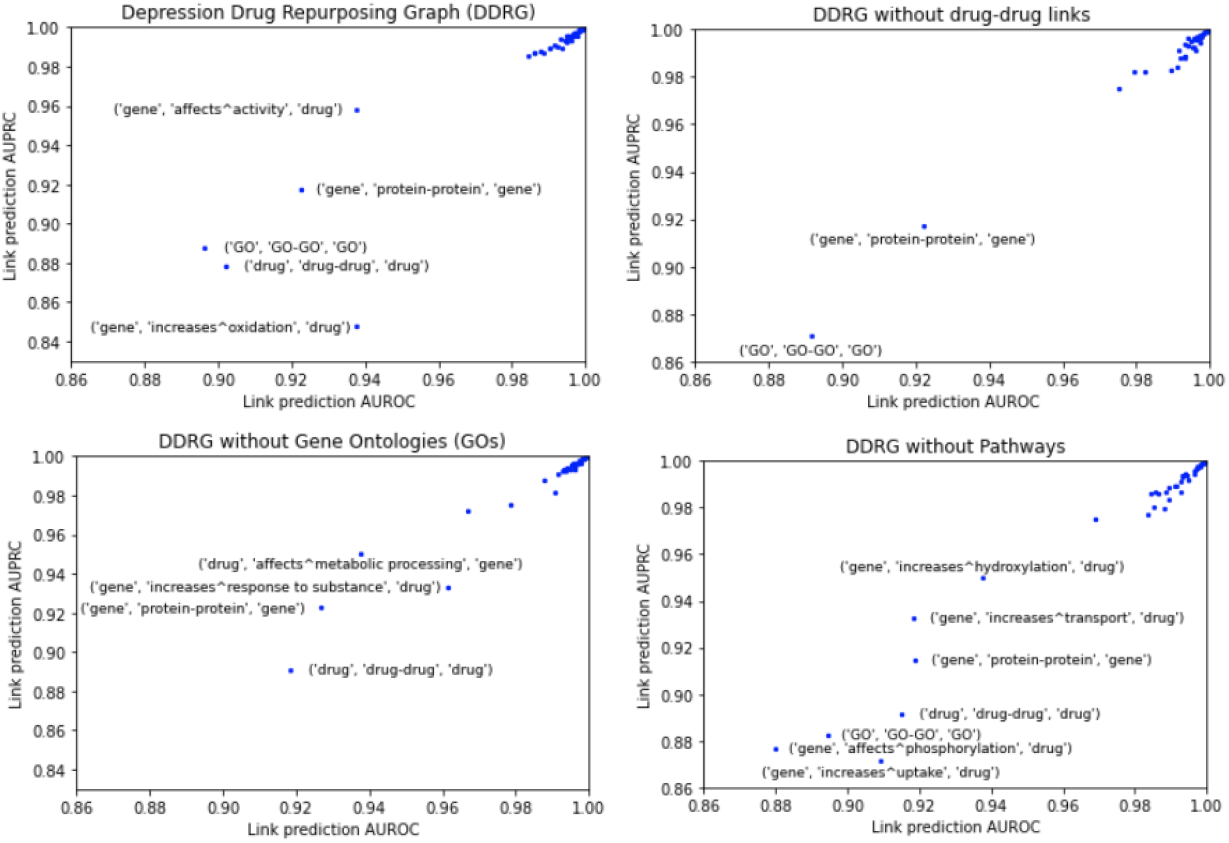
Link prediction performance at the pre-training step. Graph representation learning were evaluated by measuring link prediction performance on (A) DDRG, (B) DDRG without drug-drug edges, (C) DDRG without GO nodes, and (D) DDRG without pathway nodes.

DDRG can be denoted as an bidirectional graph *G* = (*V, E, A, R*), where *V* represents nodes, *E* represents edges, *A* represents the target and source node types, and *R* represents the edge type between the target and the source nodes. DDRG adjacency matrices *X*^*R*^ are binary matrices that indicate the edges of each edge type *R* in the graph, where 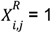 if there is an edge; otherwise, 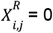.

### 5.1.1 Drug-drug similarity

The drug-drug structural similarity was computed based on four types of molecular fingerprints, including atom-pair fingerprints [36], MACCS fingerprints [37], Morgan/Circular fingerprints [38], and Topological-torsion fingerprints [39]. Sørensen–Dice similarity coefficients were calculated between drug pairs using RDkit (version 2021.03.4)[40]. Drug-drug linkages were constructed using multiple thresholds of normalized similarity coefficients *z*. Eventually, *z* ≥ 4 was selected due to the best performance.

### 5.1.2 Collection of MDD-related genes

To collect genetic markers for drug predictions, we extracted genes reported to be associated with regulations of neurotransmitters, brain neuroplasticity, neuroinflammatory mechanisms, and stress response [41]. Additionally, we extracted genes reported as either (1) diagnostic biomarkers, (2) related to disease etiology, or (3) therapeutic targets for MDD from the Comparative Toxicogenomics Database [42]. In total, 209 MDD-related genes were collected and used in the contrastive learning task (see 3.1.4).

## 6 Methods

### 6.1 Graph Representation Learning via Heterogeneous Graph Transformer (HGT)

To meaningfully transform the interaction feature of each node (drugs, genes, pathways, and GOs) in DDRG, graph representation learning is a graph embedding methodologies that embed network structures into lower-dimensional representations for the downstream task. Here, we proposed a multi-task graph representation model based on the HGT framework [31], to learn the node embeddings in DDRG. The objective is to jointly learn essential molecular interaction topology and disease genetic network for drug prediction tasks. The model includes an HGT encoder and a TransE decoder (see **Figure 1b**). The HGT parameterizes the weight matrices for the mutual attention, message passing, and propagation steps, significantly reducing required GPU memory and facilitating efficient training in heterogeneous graphs, compared with other heterogeneous GNNs.

To achieve better graph representation learning performance in sparse networks, transfer learning techniques from large-scale biomedical networks have been demonstrated to be applicable [19,22]. For example, Drug Repurposing Knowledge Graph (DRKG) [43] is an integrative biological knowledge graph that combines over five million biological interactions from multiple databases, including DrugBank, Hetionet, GNBR, String, IntAct, and DGIdb. In this study, the DRKG pre-trained embeddings of drugs, genes, GOs, and pathways in DDRG were used as initial features for graph representation learning. Embeddings were randomly generated if there are no corresponding nodes. Given each edge type, the model takes the graph adjacency matrix of DDRG and generates embeddings, which will be used for drug classification task (see Section 2.5).

#### 6.1.1 Encoder

To consider edge dependency differently across edge types in DDRG, the encoder deploys two Heterogeneous Graph Transformer (HGT) layers [31], which parameterize weight matrices for heterogeneous mutual attention, message passing, and updating steps through graph representation learning.

First, assume (*v*_*s*_, *e*_*s,t*_, *v*_*t*_) is a valid edge between node *v*_*s*_ and node *v*_*t*_ in DDRG. For the *i*^*th*^ head in the *l*^*th*^ layer, the node embedding *H*^(*l*)^ [*v*_*t*_] of node *v*_*t*_ is updated from *H*^(*l*−1)^ [*v*_*t*_] and *H*^(*l*−1)^ [*v*_*s*_]. Three linear projection functions are applied to map node embeddings into the *h*^*th*^ vector:

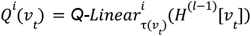

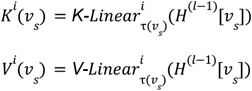

For each edge type *e*_*s,t*_ between each source and target node pairs (*s, t*), the *heterogeneous mutual attention* mechanism considers different distribution of each edge type by calculating mutual attention heads:

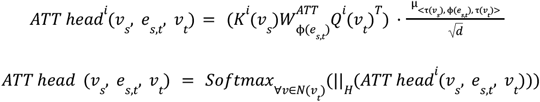

where 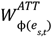 is a transformation matrix to capture edge features, T is the transposal function, and μ is an adaptive scaling to the attention of each edge (*v*_*s*_, *e*_*s,t*_, *v*_*t*_). Finally, all attention heads were concatenated and passed to a Softmax function to derive the attention head for the node pair (*v*_*s*_, *e*_*s,t*_, *v*_*t*_).

Second, the *heterogeneous message passing* mechanism extracts messages from the source node *v*_*s*_ that could be passed to *v*_*t*_ by mapping messages to multi-head attention heads:

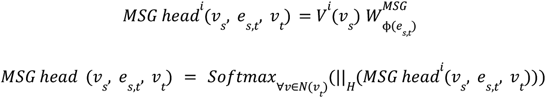

Third, the *target-specific aggregation* step updates the embedding *H*^(*l*)^ [*v*_*t*_] through a linear projection of the updated vector 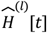, followed by a non-linear activation and residual connection:

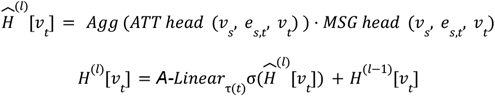

#### 6.1.2 Decoder

The encoder maps each node to an embedding that captures its connections with other nodes in DDRG. Next, a TransE decoder is used as a scoring function to measure the relationships between a source node, a target node, and their, and their relation type.

The learning objective is to minimize the distance for valid edges (true connections) while ensuring that there is a margin between the scores of valid edges and invalid edges (false or non-existent connections). This process helps encode the edge information into the learned embeddings effectively.

For each source *s* and target node *t*, and its edge type *r*, the TransE score is calculated as:

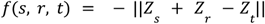

Where *Z*_*s*_, *Z*_*r*_, and *Z*_*t*_ denotes the embeddings for node *s*, relation *r*, and node *t*, respectively. *Z*_*r*_ was randomly initialized in values of [−1, 1] with the same dimension size. This scoring function ensures that embeddings *Z*_*s*_ + *Z*_*r*_ and *Z*_*t*_ are close for valid edges, and sufficiently far apart for invalid edges, effectively encoding the structure of the graph.

#### 6.1.3 Linkage prediction task

The first objective of our model is to rank positive (existing) edges over negative (non-existing) ones to learn multi-relational edge information in DDRG. With the TransE decoder, the model generates an edge score (ES) that represents the probability of existing edge between each node pair. The purpose is to learn the edge scores that capture all the edge information between existing node pairs in DDRG. For each edge type, we independently computed margin ranking loss on randomly sampled pairs of edges (*x*1, *x*2):

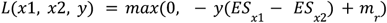

Where *y* = 0 when *x*1, *x*2 is a pair of positive edges, while *y* = 1 when *x*1, *x*2 is a pair of positive and negative edges. *ES*_*x*1_, *ES*_*x*2_ denotes the edge scores for nodes *x*1, *x*2. Additionally, *m*_*r*_ denotes a margin where larger values encourages separation between the scores of positive and negative edges.

#### 6.1.4 Contrastive learning task

The second objective of our model is to learn similar embeddings for known or potential disease targets, disease-causing genes, and genetic predispositions. Since positive drugs often share targets, this step could help classify between positive drugs and negative drugs. Contrastive learning is a self-supervised learning technique that aims to derive similar representations for samples in the same class while generating dissimilar embeddings for samples in distinct classes. Here, a contrastive learning task was used to simultaneously learn MDD gene class information to distinguish between depression-related genes and the rest. We computed contrastive loss for each pair of genes (*g*1, *g*2):

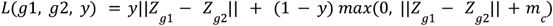

Where *y* = 0 when *g*1, *g*2 is a pair of positive genes, while *y* = 1 when *g*1, *g*2 is a pair of positive and negative genes. *Z*_*g*1_, *Z*_*g*2_ denotes the embeddings for genes g1 and g2. The loss minimizes the distance for positive pairs and maximizes it for negative pairs beyond the margin *m*_*c*_.

#### 6.1.5 Model training and evaluation

As described above, our multi-task representation learning model has two objective functions in the optimization process. To avoid conflict between the multiple objective functions, we used a gradient surgery library called PcGrad [44], which projects the conflicting gradient into the other objective’s gradient. PcGrad was implemented in PyTorch to achieve a better local optimum.

### 6.2 Drug Classification Model via Relational Graph Convolutional Network

The drug classification task was also implemented using the Relational Graph Convolutional Network [30]. 851 (70%) of all 1,216 drug candidates were used to train the classification model. Among all drugs, 63 approved or investigational medications for MDD or have reported therapeutic roles on MDD were collected from Drug Repurposing Hub [45], DrugBank [46], and Comparative Toxicogenomics Database [42], which were used as positive drugs, while the rest of the drugs were set as negative labels.

In the drug classification task, the DDRG drug embeddings learned by the graph representation model were input into the drug classification model, which predicts drug classes. To deal with the imbalance issue of class labels, we utilized focal loss for model training:

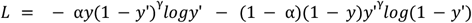

Where modulating factors α and focusing parameters *γ* are tunable parameters that aim to focus learning on hard misclassified samples. For instance, α closer to 1 places greater emphasis on positive cases (*y* = 1). If *γ* > 0, the function downweights the easy examples where *y*′ is close to the true label *y*.

## Data availability

All data and code in the study are available on a GitHub repository at https://github.com/khlin65/HeteroRepur.

